# Comparisons between Complete Genomes of the Eukaryotic Extremophile *Galdieria sulphuraria* Reveals Complex Nuclear Chromosomal Structures

**DOI:** 10.1101/2022.10.04.510839

**Authors:** Jessica M. Downing, Sarah C. L. Lock, Manuela Iovinella, John Davey, Luke C. M. Mackinder, James P. J. Chong, Peter D. Ashton, Georg A. Feichtinger, Sally James, Daniel Jeffares, Claudia Ciniglia, Seth J. Davis

## Abstract

Extremophiles, while typically bacteria and archaea, are also found in the eukaryotic domain of life. The eukaryote *Galdieria sulphuraria* is a thermoacidophilic red alga belonging to the class *Cyanidiophyceae*, an especially unique class as it comprises the basal clade of eukaryotic extremophiles. *Galdieria* species can grow both photosynthetically and heterotrophically on a variety of carbon sources, thriving down to pH 0 and temperatures up to 56 °C, while tolerating high levels of reactive oxygen species and high levels of heavy metals. Here we report whole-genome sequencing of three *G. sulphuraria* strains, uncovering a compact (13.1 Mb – 16.0 Mb) nuclear genome with 72-73 chromosomes, dependent on the strain.Comparative analyses of the macro synteny revealed significant structural rearrangement between *G. sulphuraria* isolates and the genome shows signs of sexual recombination. This, along with the large number of nuclear chromosomes compared to the genome size, reveals a mechanism of intrinsic adaptability in this eukaryotic extremophile, uncovering how *G. sulphuraria* can thrive in a rapidly changing extreme environment.

---

Extremophiles are organisms that are adapted to life in extreme conditions including extreme temperatures (e.g. <10 °C, >50 °C), extreme pH (e.g. <3, >10), hypersaline environments, high pressure, and harmful radiation levels [1]. They are of great interest since extreme conditions were widespread on early earth, therefore extremophiles are useful models to study for understanding early life [2]. Moreover, extremophiles are a source of novel enzymes that are tolerant to extreme conditions and have applied uses in industry [3]–[6], most notably Taq polymerase, isolated from the thermotolerant eubacteria, *Thermus aquaticus* [7], which revolutionised molecular biology. Extremophiles are found in all kingdoms of life.

An example of known photosynthetic extremophiles belong to the *Cyanidiophyceae*, a class of unicellular red algae. They are thought to have diverged from the rest of the red algal lineage around 1.3 billion years ago, the class contains four genera, the mesophilic *Cyanidium*, and the thermoacidophilic *Cyanindioschyzon, Cyanidiococcus*, and *Galdieria*, and at least six species [8]. To date, only one *Cyanidiophyceae* genome has been completely resolved. *Cyanidioschyzon merolae* has a nuclear genome size of 16.5Mb and 20 chromosomes. The plastid and mitochondrial genomes have also been resolved and are 150,000bp and 32,200bp in size respectively [9].

*Galdieria sulphuraria* is an extremely successful member of the *Cyanidiophyceae*. It thrives at relatively high temperatures (up to 56 °C), and low pH (pH 0-4) [10]. Additionally, *G. sulphuraria* tolerates high levels of heavy metals and reactive oxygen species, growing in both hot springs, and acid mine drainage sites. *Galdieria* species are the only known members of the *cyandiophyceae* that are not obligately phototrophic. In addition to using light as a fuel source, *G. sulphuraria* utilises a variety of complex carbon sources for growth [11]. Consequently, *G. sulphuraria* is of great interest for use in biotechnology for the degradation of lignocellulosic materials [12], as well as the remediation of rare earth metals [13][14] and wastewater treatments [15].

Whole genome sequencing experiments have revealed that, like *C. merolae, G. sulphuraria* has a compact genome, between 12Mb and 15Mb in size. This is typical of a Rhodophyte since Rhodophyte genomes are compact, with between ∼5000-10000 genes. This has been attributed this to two phases of significant gene loss in the red algal common ancestor, with an initial genome reduction resulting in the loss of around 25% of its ancestral gene complement [16]. Analyses of the *Galdieria* genomes reveal that around 1% of *G. sulphuraria* genes were obtained through horizontal gene transfer from neighbouring bacteria and archaea [17]. These HGT genes include genes which may confer *G. sulphuraria* resistance to specific extreme conditions and may have been an evolutionary driver for extremophilicity in *G. sulphuraria* [18].

High quality genome data would be particularly useful for understanding *Galdieria* biology. In the absence of a pre-existing reference genome, 2^nd^ generation short read sequencing faces a major challenge in the construction of long contigs. There have been attempts to quantify the number of chromosomes in *G. sulphuraria* using pulse field gel electrophoresis, estimating a genome size of 9.8 Mb and 40 chromosomes [19], indicating that *G. sulphuraria* possesses many chromosomes considering the small genome size. The 2016 *G. sulphuraria* SAG 107.79 assembly resolved into 117 contigs with a contig N50 of 134Kb [20]. Although this assembly was incomplete, it provided a useful example of how Oxford Nanopore Sequencing can be applied to the assembly of whole genomes. The NCBI reference genome for *G. sulphuraria* 074, originally isolated from Java, Indonesia, sequenced using Sanger and 454 sequencing, is assembled into 433 contigs. The construction of complete *Galdieria* genomes is critically important for the examination of how chromosomal architecture supports the adaptive, flexible, and versatile life capacity of this species. Here we describe the reassembly of *G. sulphuraria* SAG 107.79, as well as the sequencing and assembly of 2 further *G. sulphuraria* strains, ACUF 138 and ACUF 017 into complete genomes.

## Methods

### Strain Preparation

*G. sulphuraria* ACUF 138 and ACUF 017, originally isolated from El Salvador and the Phlegraean Fields, Italy, respectively, were obtained from the Algal Collection of University of Naples (ACUF) [21]. *G. sulphuraria* SAG 107.79, originally isolated from Yellowstone National Park, USA, was obtained from the Culture Collection of Algae (SAG) at Göttingen University [22]. All strains were isolated from a single colony obtained after streaking the culture across agar plates, respectively, and colonies were inoculated in Allen medium pH 1.5 [23] and cultivated at 37°C under continuous fluorescent illumination of 45 µmol photons·m^−2^·s^−1^.

### DNA Extraction

DNA extraction protocols are detailed in Supplementary Materials 1.

### DNA Sequencing

Library preparations were conducted using NEBNext® Ultra™ II DNA Library Prep Kit for Illumina Sequencing according to the manufacturer ‘s instructions. Libraries were then sequenced with Illumina MiSeq (Illumina, San Diego, CA) and the resulted reads were trimmed with Trimmomatic [24] and assembled using Spades v3.1 [25].

### Assembly

Oxford Nanopore MinION reads were base called with guppy 4.0.11 [26] with options --flowcell FLO-MIN106, --kit SQK-LSK108, --trim_strategy dna and --trim_barcodes.

Three draft assemblies were generated from each set of nanopore reads with the assemblers Canu2.1 [27], Raven v1.5.3 [28], and SMARTdenovo [29]. Canu2.1 was run with options genomeSize=13m and -fast. SMARTdenovo was run in consensus mode, -c 1.

Each draft assembly was polished once with medaka v1.3.3 [30], and polished three times with Pilon v1.23 [31], using the Illumina reads mapped to the assembly using the Burrows-Wheel Aligner v0.7.17 [32].

Assemblies were assessed with Tapestry v1.0.0 [33], aligned to each other by minimap2 v2.20 [34], and edited in Biopython. For *G. sulphuraria* ACUF 138, contigs with less than 10% unique material were removed from each polished assembly. The final assembly was compiled from contigs from the Canu2.1 and SMARTdenovo assemblies by manually checking each contig pair for which contig had telomeres, and checking the read alignments for gaps that could suggest a misassembly. The final assemblies were checked with Tapestry and annotated as described below.

To assess genome structural diversity between strains, assemblies were aligned to one another by nucmer (MUMmer v 4.0.0). *G. sulphuraria* SAG 107.79 was aligned to *C. merolae* 10D by promer [35]. Dotplots were visualised with gnuplot v 5.2.8 [36]. Assemblies were visualised using RStudio KaryoplotR.

### RNA Preparation and Extraction

Protocols for RNA preparation and extraction are detailed in Supplementary Materials 2.

### Genome Annotation

Transcript assemblies were constructed using Trinity using both de-novo and genome guided modes [37]. RNA sequencing reads to their respective Illumina assemblies using the splice aware aligner STAR v. 2.7.3 [38]. Annotations were predicted with funannotate [39], run without coding quarry, using the eukaryota database (fetched 02/03/2021) for functional predictions [40]. InterProScan v 5.46-81.0 [41] was run separately.

### Variant Calling and Linkage Disequilibrium

Illumina reads from the entire collection of *G. sulphuraria* isolates described in the companion paper (Lock+Iovinella et al. 2022). These were aligned to the completed *G. sulphuraria* SAG 107.79 genome using the Burrow-Wheeler Aligner v.0.7.17 [32]. Aligned reads were processed using SAMtools v.1.10 [42] and the Picard Toolkit v 2.21.6 MarkDuplicates and AddOrReplaceReadGroups [43]. Variants were called using the Genome Analysis Toolkit v.4.1.0.0 (GATK) [44]. Repetitive, difficult to map regions were excluded from the analysis. These were identified as regions with high depth of coverage as determined by mosdepth v.0.2.8 [45] after all by all chromosome alignments using minimap2 [34] in mode -ax ava-ont. Hard filtering was applied using BCFtools v.1.10.2 [46] excluding sites with an RMS Mapping Quality (MQ), a Phred-Score (FS), Quality by Depth (QD) of < 32. The linkage disequilibrium correlation co-efficient (R^2^) values were calculated using PLINK v.2.00 [47], [48]. After calculating the mean R^2^ value over 1Kb pairwise distance windows, Spearman ‘s correlation coefficient was calculated using RStudio.

## Results

The completed genome assemblies for *G. sulphuraria* SAG 107.79, ACUF 138 and ACUF 017 reveal a highly compact genome with an unusually large number of chromosomes compared with the genome size. The genome size ranged from 13 Mb to 16 Mb and the number of contigs was 72-73, strain dependent. The genome size is consistent with previously reported *G. sulphuraria* genome sequences. Two strains, *G. sulphuraria* SAG 107.79 and *G. sulphuraria* ACUF 138, have complete BUSCOs > 90%, consistent with the assemblies that include the majority of genes. Of the two, *G. sulphuraria* SAG 107.79 is the most complete genome by measure of the number of telomere-to-telomere chromosomes. Although *G. sulphuraria* ACUF 138 has a higher scaffold N50 value and a slightly higher % Complete BUSCOs, these differences can be attributed to the larger genome size of *G. sulphuraria* ACUF 138, at 15.95 Mbp compared to 14.28 Mbp in *G. sulphuraria* SAG 107.79.

**Table 1.** Assembly and annotation statistics for G. sulphuraria strains SAG 107.79, ACUF 138 and ACUF 017, alongside the closest available complete genome, Cyanidioschyszon merolae, and the previously published G. sulphuraria reference genome, 074W. ^*^Assuming a 13Mb genome

|  | *G. sulphuraria* Assemblies |  |  | Closest Complete Genome | *G. sulphuraria* NCBI |
| --- | --- | --- | --- | --- | --- |
|  | ACUF 138 | SAG 107.79 | ACUF 017 | *Cyanidioschyzon merolae* 10D | 074W |
| Assembly Size (Mbp) | 15.95 | 14.28 | 13.14 | 16.52 | 13.78 |
| Largest Scaffold (bp) | 385618 | 500147 | 349111 | 1621617 | N/A |
| Average Scaffold (bp) | 218495 | 198347 | 182544 | 826015 | 31824 |
| Num Scaffolds | 73 | 72 | 72 | 20 | 433 |
| Scaffold N50 (bp) | 221800 | 209122 | 196143 | 850100 | 172100 |
| % GC | 39.04 | 40.19 | 39.06 | 55.00 | 36.89 |
| Num Genes | 6404 | 5975 | 5567 | 5331 | 6723 |
| Num Proteins | 6357 | 5920 | 5523 | 5010 | 6622 |
| Num tRNA | 47 | 55 | 44 | 30 | 127 |
| % Genes of Unknown Function | 21.9 | 19.62 | 20.94 | N/A | N/A |
| Gene Density | 2491 | 2390 | 2361 | 3099 | 2050 |
| Introns Per Gene | 2.83 | 2.93 | 2.92 | 0.005 | 1.26 |
| Mean Gene length | 1592 | 1598 | 1571 | 1552 | N/A |
| % Coding | 54.99 | 57.66 | 57.34 | 44.9 | N/A |
| % Complete BUSCOs | 90.5 | 90.1 | 88.5 | N/A | N/A |
| Contigs Telomere-Telomere | 50 | 62 | 18 | N/A | N/A |
| **Contigs with 1 Telomere** | 20 | 10 | 38 | N/A | N/A |
| **Contigs with No Telomeres** | 3 | 0 | 16 | N/A | N/A |
| **ONT Read Coverage > 15Kb\*** | 455 | 220 | 13 | N/A | N/A |

The *G. sulphuraria* ACUF 017 assembly has fewer telomere-to-telomere chromosomes, a lower N50 value and Complete BUSCOs < 90%. This is due to lower quality raw sequencing data for this sample, with low coverage of ONT reads longer than 15 Kb that would be required to bridge over the numerous subtelomeric regions in the *G. sulphuraria* genome assembly. Comparisons of the *G. sulphuraria* SAG 107.79 and *G. sulphuraria* ACUF 138 assemblies reveal important structural differences. *G. sulphuraria* ACUF 138 chromosome length is more uniform than that of *G. sulphuraria* SAG 107.79. The longest contig in *G. sulphuraria* ACUF 138 is 385.6 Kb, and the shortest is 131.4 Kb. *G. sulphuraria* SAG 107.79 has a longer longest contig, SAG 107.79:scaffold_1, at 500.1 Kb, and a shorter shortest contig at 62.6 Kb, although this contig is missing a telomere. The shortest telomere to telomere contig for *G. sulphuraria* SAG 107.79 is 84.8 Kb. SAG 107.79:scaffold_1 is longer than the next longest chromosome by 144.0 Kb.

Alignments of *G. sulphuraria* SAG 107.79 vs *G. sulphuraria* ACUF 138 show that SAG 107.79:scaffold_1, the longest chromosome of *G. sulphuraria* SAG 107.79, aligns to two chromosomes in *G. sulphuraria* ACUF 138, ACUF 138:scaffold_3 and ACUF 138:scaffold_14, revealing that SAG 107.79:scaffold_1 is either the result of a fusion between two chromosomes, or has broken into two chromosomes, at some point in the lineage, explaining why *G. sulphuraria* SAG 107.79 has a longest chromosome 114.5 Kb longer than ACUF 138:scaffold_1. Other structural variations are observed between these two genomes, with SAG 107.79:scaffold_5 aligning to ACUF 138:scaffold_7 and ACUF 138:scaffold_10, and SAG 107.79:scaffold_8 aligning to ACUF 138:scaffold_38 and ACUF 138:scaffold_40. These structural rearrangements are unidirectional – there were no instances found of *G. sulphuraria* ACUF 138 chromosomes being “split” into two chromosomes in *G. sulphuraria* SAG 107.79 (figure 2).

**Figure 1:**
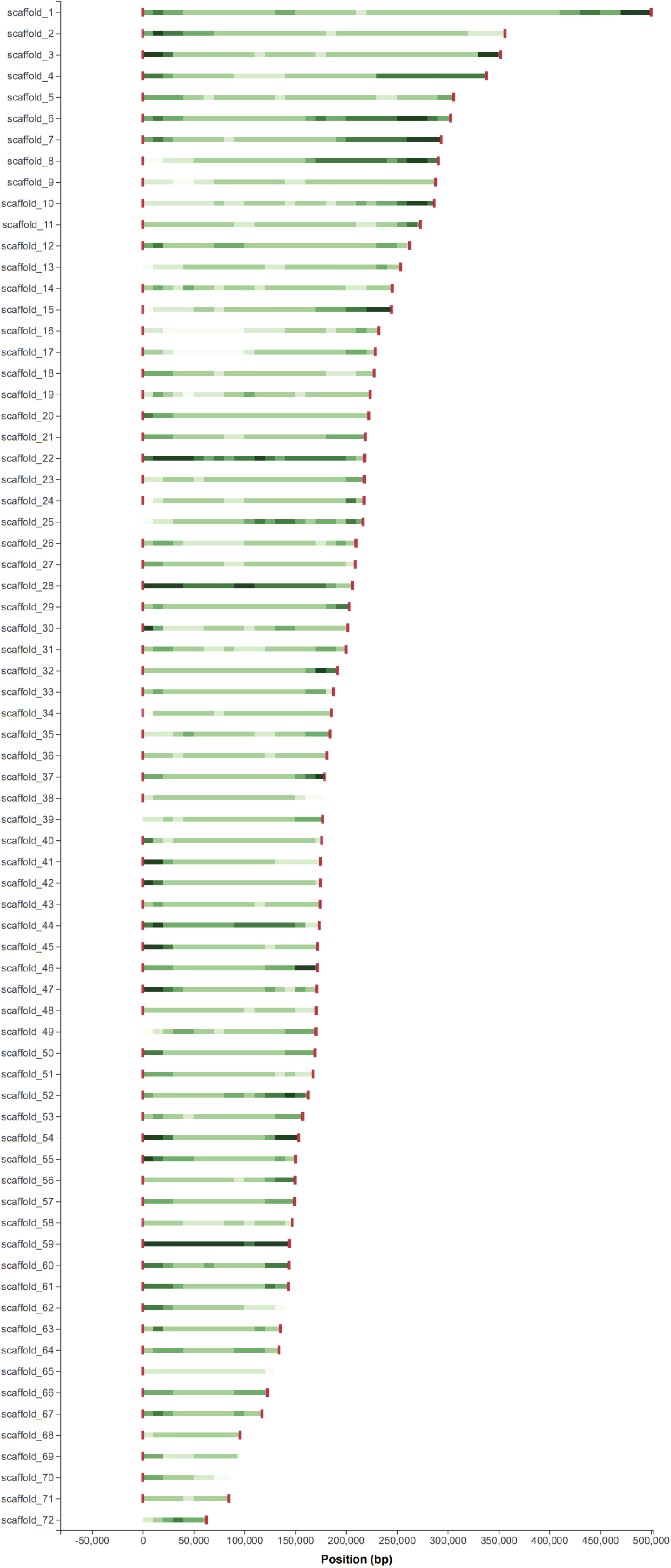
Complete G. sulphuraria SAG 107.79 nuclear genome assembly. For more detailed figures of G. sulphuraria SAG 107.79, and G. sulphuraria ACUF 138, see Supplementary Figures 1+2.

**Figure 2:**
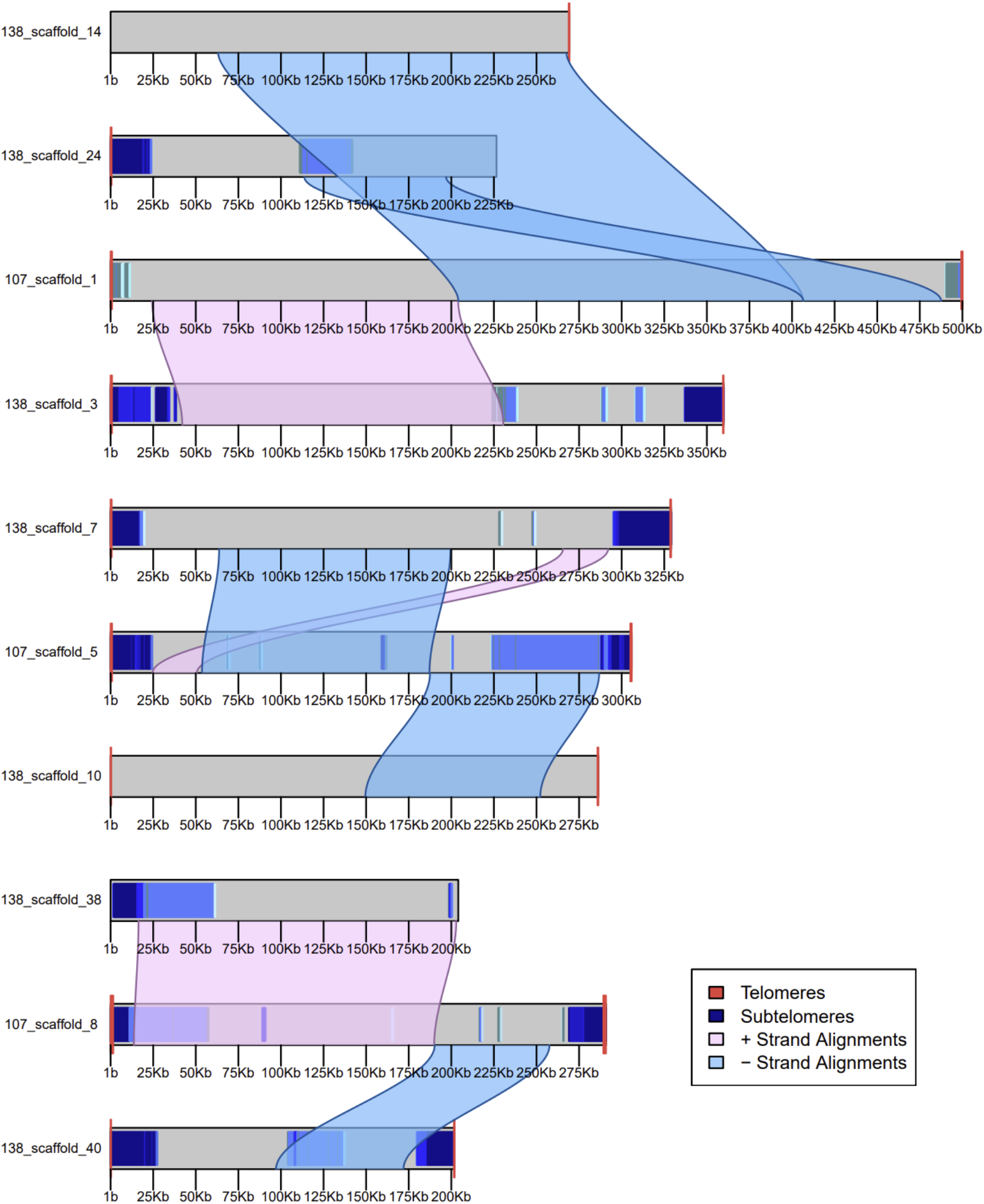
Alignments of G. sulphuraria SAG 107.79 scaffolds 1, 5, and 8 to G. sulphuraria ACUF 138, generated by AnchorWave [49]. For whole genome alignments, see Supplementary Materials 3.

Similar rearrangements are seen in the alignment between *G. sulphuraria* SAG 107.79 and *G. sulphuraria* ACUF 017. ACUF 017:scaffold_5, which aligns to SAG 107.79:scaffold_1 along with ACUF 017:scaffold_6 and ACUF 017:scaffold_24, is missing one telomere, but contains subtelomeric repeats, higher coverage depth and lower gene density at the end of the contig, indicating this is a complete chromosome and that SAG 107.79:scaffold_1 is divided into at least two chromosomes in *G. sulphuraria* ACUF 017 in addition to *G. sulphuraria* ACUF 138. Chromosome rearrangements are not unidirectional between ACUF 017 and SAG 107.79, with ACUF 017:scaffold_28 aligning to 5 *G. sulphuraria* SAG 107.79 chromosomes (Supplementary Materials 3).

To investigate the possibility that recombination is a driving factor in some of these structural changes, linkage disequilibrium (LD) was measured for intra- and inter-chromosomal SNPs. For intrachromosomal SNPs, LD was observed to decay with pairwise distance (figure 3), consistent with meiotic recombination.

**Figure 3:**
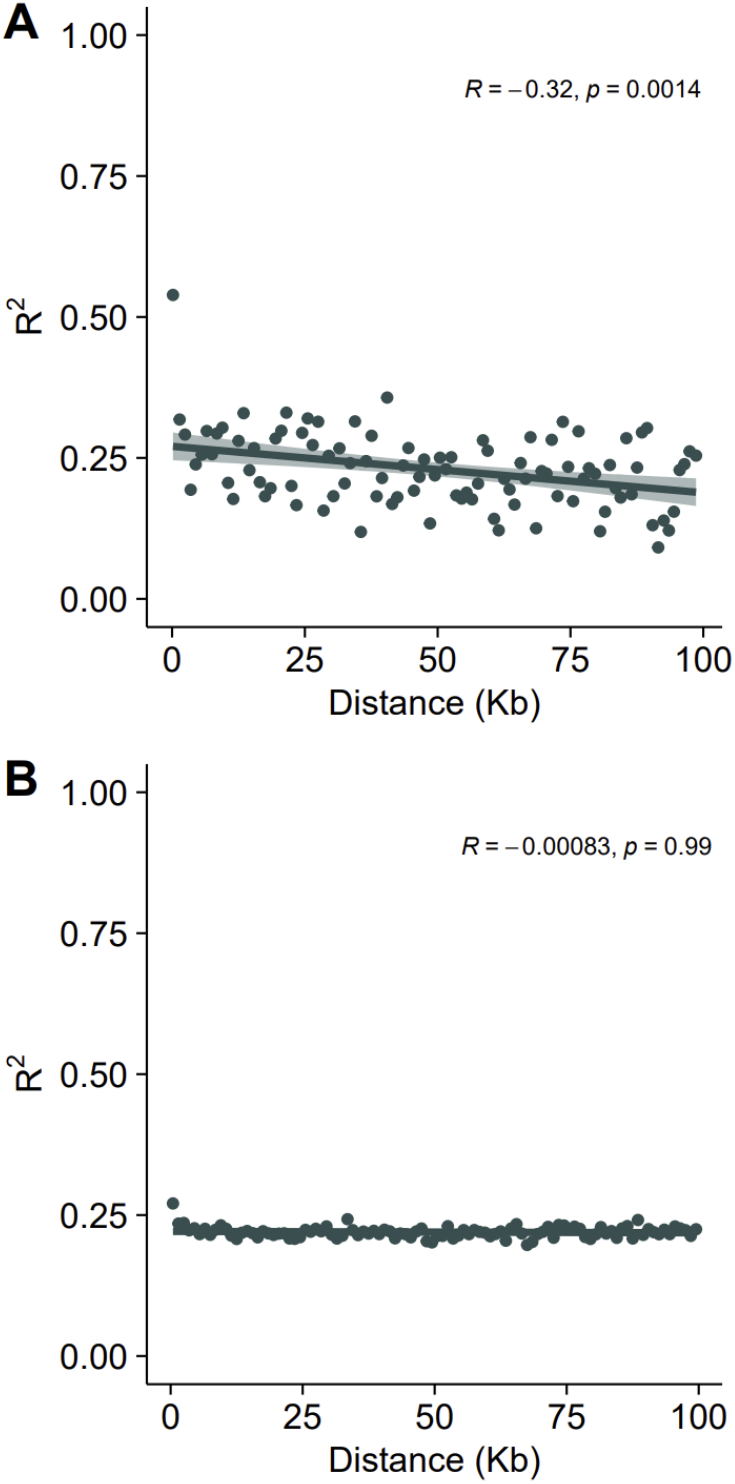
Mean linkage disequilibrium coefficient, R^2^ against mean pairwise distance in 1Kb intervals between a) SNPs on the same chromosome and b) all SNPs. Spearman ‘s R calculated using ggpubr.

## Discussion

Here we have described the first complete genomes of the polyextremophile*G. sulphuraria*, revealing a highly unusual genome structure – with an estimated 72 chromosomes for a genome size of 13.14 Mb (*G. sulphuraria* SAG 107.79). While it is not uncommon to find species with numbers of chromosomes in excess of 70, considering the relatively small genome size of *G. sulphuraria*, the number of nuclear chromosomes is especially high. Other small eukaryote genomes such as that of *Galdieria* ‘s relative *Cyanidioschyzon merolae* 10D (16.52 Mb) [9], and the green alga *Ostreococcus tauri* (12.56 Mb) [50] have much lower numbers of chromosomes – 20 for both species. This number of chromosomes is significantly higher than previously estimated by Moreira et al. [19], however this experiment underestimated the size of the *G. sulphuraria* genome by at least 3 Mb. This is likely because *G. sulphuraria* contains a lot of small chromosomes that would be indistinguishable on a pulse field gel. Here we show that *G. sulphuraria* has multiple chromosomes within the 100 Kb to 200 Kb size range, had all of these been attributed to a single or a few gel bands, this would cover the underestimation in genome size and number of chromosomes. Additionally, the number of genes and genome size of these assemblies is consistent with the previously available *G. sulphuraria* 074W genome [18]. Why *G. sulphuraria* has this many chromosomes is unclear, but since *G. sulphuraria* is extremely dominant in its environment [10], it does raise questions as to whether having many small chromosomes (as opposed to fewer, larger, chromosomes) could provide an evolutionary advantage for eukaryotes in extreme environments, by allowing for increased adaptation.

The most complete *G. sulphuraria* is *G. sulphuraria* SAG 107.79, since it has the most complete (telomere-to-telomere) chromosomes and has a BUSCO score > 90%. *G. sulphuraria* ACUF 138 is as complete as *G. sulphuraria* SAG 107.79 in terms of complete BUSCOs, but is missing more telomeric regions, however since the telomeres tend not to harbor coding material, this is of little consequence. *G. sulphuraria* ACUF 017 is less complete, and would benefit from resequencing with gentler DNA extraction methods to achieve higher coverage depth of longer reads. Nevertheless, the *G. sulphuraria* ACUF 017 assembly is consistent in size and chromosome number to the *G. sulphuraria* SAG 107.79 and ACUF 138 assemblies, which were constructed from higher quality raw sequencing data.

The next question surrounds the mechanism driving the chromosomal structural diversity between *G. sulphuraria* strains, as well as the duplications/shared regions between chromosomes in a single assembly (Supplementary Materials 4). As previously mentioned, *G. sulphuraria* SAG 107.79 has a longest chromosome 114.5 Kb longer than that of *G. sulphuraria* ACUF 138, and 151.0 Kb longer than the longest chromosome of *G. sulphuraria* ACUF 017. Interestingly, attempting to use the *G. sulphuraria* SAG 107.79 assembly to train medaka to polish the *G. sulphuraria* ACUF 017 SMARTdenovo assembly fails (data not shown), indicating that the two genomes are too different to use for scaffolding. In addition to the differences in genome structure, phylogenomic data from Lock + Iovinella demonstrates that the pangenomes of these three strains are more distant than one would expect for different strains from the same species. Given the karyotypic and sequence diversity between these strains, it is not implausible that these are different species, however since they are not morphologically distinct, this claim needs to be investigated further.

It has previously been reported that *G. sulphuraria* genomes contain the molecular machinery to facilitate meiotic recombination [51]. Although the *Cyanidiophyceae* was assumed to be asexual, there is increasing evidence that sexual reproduction is ubiquitous among eukaryotes, and assumed asexuality is as a result of lack of study and observance of these processes [52]–[55]. Illumina read alignments show that the *G. sulphuraria* genome is diploid, with numerous heterozygous sites, and *G. sulphuraria* SAG 107.79 Illumina reads achieved by resequencing the genome after propagation of a single colony over a five-month period revealed that recombination events have taken place followed by haplotype loss (Supplementary Materials 5). The presence of linkage disequilibrium is also consistent with meiotic recombination. Finally, the nuclear, mitochondrial, and plastid phylogenies are incongruent (Lock + Iovinella). Taken together, these data comprise strong molecular evidence that *G. sulphuraria* is a diploid organism that undergoes sexual recombination.

Recombination has traditionally been theorized to confer an evolutionary advantage by creating increased genetic diversity in populations [56], and has been shown to be a driving factor in enabling parasites to adapt to environmental conditions and evade host responses [57]. For *G. sulphuraria*, an organism that must adapt to a rapidly changing extreme environment, recombination could serve a similar purpose, however the exact mechanism by which *G. sulphuraria* may conduct sex, is unknown. Efforts are ongoing to fully phase the *G. sulphuraria* genome.

The complete genome of *G. sulphuraria* has revealed secrets on some of the potential molecular mechanisms of extreme adaptation in this eukaryote, demonstrating that the mechanisms for extremophilicity in this eukaryote are multi-faceted and not only limited to previously established mechanisms such as horizontal gene transfer [18]. With the benefit of a complete *G. sulphuraria* genome, investigations can continue into furthering the understanding of the inner workings of extremophiles.

## Notes

### Competing Interest Statement

The authors have declared no competing interest.

