## Supplementary for "Comparisons between Complete Genomes of the Eukaryotic Extremophile *Galdieria sulphuraria* Reveals Complex Nuclear Chromosomal Structures"

### Supplementary Materials

#### 1. DNA Extraction

Illumina Sequencing

After a culture was grown to stationary phase 2 ml of a given culture was centrifuged (5 minutes, 13.2 rpm) and the algal pellet was resuspended in 40 µl PBS pH 7.5 and vortexed to mix. The samples were centrifuged and resuspended in PBS a further two times. Tubes were placed in a dry ice ethanol bath for 30 seconds then transferred to a 30˚C water bath, this was repeated 3 times.
From there, 500 µl of DNA extraction buffer 1 was added and incubated for 30 minutes at 55 ˚C, inverting every 10 minutes. Then 150 µl of DNA extraction buffer 2 was added and the samples incubated for a further 10 minutes at 65 ˚C. Finally, 650 µl of phenol:chloroform:isoamyl alcohol 25:24:1 was added. To this, 1 mm silica beads were added (0.5 cm of the tube), the samples were mixed by inversion, and mixed on a bead beating machine for 5 minutes. After centrifugation to clarify this solution, 600 µl of the supernatant was taken into a fresh tube and 480 µl of cold isopropanol was added. The samples were stored at -20 ˚C for a minimum of 2 hours. Next, samples were centrifuged at 15000 g for 30 minutes at 4 ˚C, and the supernatant discarded. 200 µl of ethanol was added then centrifuged at 13.2 rpm for 5 minutes and the supernatant discarded. The tubes were dried at room temperature, then resuspended in 30 µl of Tris-EDTA.

Samples were incubated with 1 µl of RNAse A and 1 µl of Proteinase K for 2 hours at 37 ˚C, then purified using the Qiagen DNeasy Plant Mini Kit. DNA quality and concentration were assessing using a Nanodrop photospectrometer ND-1000 (Thermo Fisher Scientific).

Nanopore Sequencing

After a culture was grown to stationary phase, 2 ml of a given culture was centrifuged for 5 minutes at 13.2rpm and supernatant discarded. Tubes were placed in a dry ice ethanol bath for 30 seconds then transferred to a 30˚C water bath, this was repeated 4 times. Next 1µl of Protienase K, 100µl of Viscozyme™ were added and incubated for an  hour at 37˚C. Then 40µl of PBS pH 7.5 added and vortexed to mix. 500µl of DNA extraction buffer 1.1 was added and incubated at 55˚C for 30 minutes, mixing by inverting every 10 minutes. Then 150µl of DNA extraction buffer 2 (2.1 or 2.2 dependent on strain) was added and incubated for a further 10 minutes at 65˚C. Next 690µl of Phenol:Chloroform:Isoamyl Alcohol 25:24:1 was added and mixed gently though inversion for 5 minutes. This was centrifuged at 13.2rpm for 5 minutes and 600µl of the top layer of the supernatant was then taken and placed into a fresh tube. Here 480µl of isopropanol was added and samples stored at -20˚C for 2 hours. Following this, samples were centrifuged at 15g for 30 minutes at 4˚C, supernatant then discarded. 200µl of 70% ethanol was added and then centrifuged at 13.2rpm for 5 minutes, supernatant discarded. Finally, tubes were air dried and then DNA re-suspended in 40µl of TE buffer.

Clean-up of DNA samples were completed using Zymo DNA Clean & Concentrator™-25 kit, the method was as follows: DNA binding buffer (DNA Binding Buffer: DNA sample) were added to DNA samples in a ratio of 2:1 and mixed briefly by vortexing. The mixture was transferred to a Zymo-Spin™ Column in a Collection Tube and was centrifuged for 30 seconds at 13.2rpm. The flow-through was discarded. 200µl DNA Wash Buffer was added to the column and centrifuged for 30 seconds at 13.2 rpm and the flow-through discarded. This wash step was repeated. It was then again centrifuged at 13.2rpm for 30 seconds and 40µl DNA Elution Buffer at 65˚C was slowly added directly to the column matrix and incubated at room temperature for ten minutes. The column was then transferred to a 1.5ml microcentrifuge tube and centrifuged at for 30 seconds at 13.2rpm to elute the DNA. This step was repeated once. DNA quality and concentration were assessing using a Nanodrop photospectrometer ND-1000 (Thermo Fisher Scientific).

| **DNA Extraction buffers** | | |
|---|---|---|
| **Buffer 1.1** | **Buffer 2.1** | **Buffer 2.2** |
| 200mM Tris-HCl pH8 | 200mM Tris-HCl pH8 | 100mM Tris-HCl pH8 |
| 200mM NaCl | 200mM NaCl | 700mM NaCl |
| 100mM LiCl | 100mM LiCl | 20mM EDTA pH8 |
| 25mM EDTA pH8 | 25mM EDTA pH8 | 2% CTAB |
| 1M Urea | 1M Urea | 0.0125mM PVP-40 |

| | |
|---|---|
| 1% SDS | 1% CTAP |
| 1% NP-40 | 100mM Lithium acetate |

*Table 1: DNA Extraction Buffers*

### 2. RNA Preparation

*G. sulphuraria* cultures (017, 033, 074, 107, 138 and 427) were grown under a 12h/12h light/dark cycle under 42 µmol $m^{-2}$ $s^{-1}$ at 37°C on an orbital shaker (130rpm). The experimental design followed different growth conditions to obtain a great variety of mRNAs. Samples were grown in Allen medium mixotrophically with 10 g/L Sucrose at pH 2, in Allen medium with 0.5% Cellulose (w/v), 0.5% Xylan (w/v) and 0.5% Laminarin (w/v) at pH 2. Samples were collected by centrifugation at 1h, 12h, 96h, 192h and 336h. Pellets were washed 3 times in PBS buffer (pH 7.5) and samples stored at -80 ˚C before RNA extraction.

Cells were ground into a fine powder with a pestle and a drill in the presence of liquid nitrogen. RNA was isolated and cleaned up using the Monarch Total RNA Miniprep Kit (New England BioLabs, T2010S). RNA quality and concentration was assessed using a Nanodrop photospectrometer ND-1000 (Thermo Fisher Scientific). All RNAs were treated with DNaseI and then pooled by strain relative to the concentration of each sample. RNA quality and concentration was then assessed using the Agilent 2100 Bioanalyzer (Agilent Technologies).
RNA library preparation and sequencing were performed at Novogene (UK) Company Limited (Cambridge). Library preparation was performed using NEB Next® Ultra™ RNA Library Prep Kit (NEB, San Diego, CA, USA), employing AMPure XP Beads to purify the products of the reactions during the library prep. Poly-a mRNA was isolated using poly-T oligo-attached magnetic beads, then fragmented through sonication and enriched into 250-300bp fragments. The purified mRNA was converted to cDNA and subjected to the adaptor ligation. The barcoded fragments were finally multiplexed and ran on the Illumina Novaseq 6000 (s4 flow cell) to acquire 20 million read pairs per sample, using the 150bp PE sequencing mode.

### 3. Alignments of G. *sulphuraria* Assemblies

Whole genome alignments of a) *G. sulphuraria* ACUF 017 to *G. sulphuraria* ACUF 138, b) *G. sulphuraria* ACUF 017 to *G. sulphuraria* SAG 107.79, and c) G. sulphuraria ACUF 138 to *G. sulphuraria* SAG 107.79. All alignments were generated in nucmer, and filtered to remove alignments <100nt [1]. Forward strand alignments are shown in purple and reverse strand alignments are shown in blue.

a)

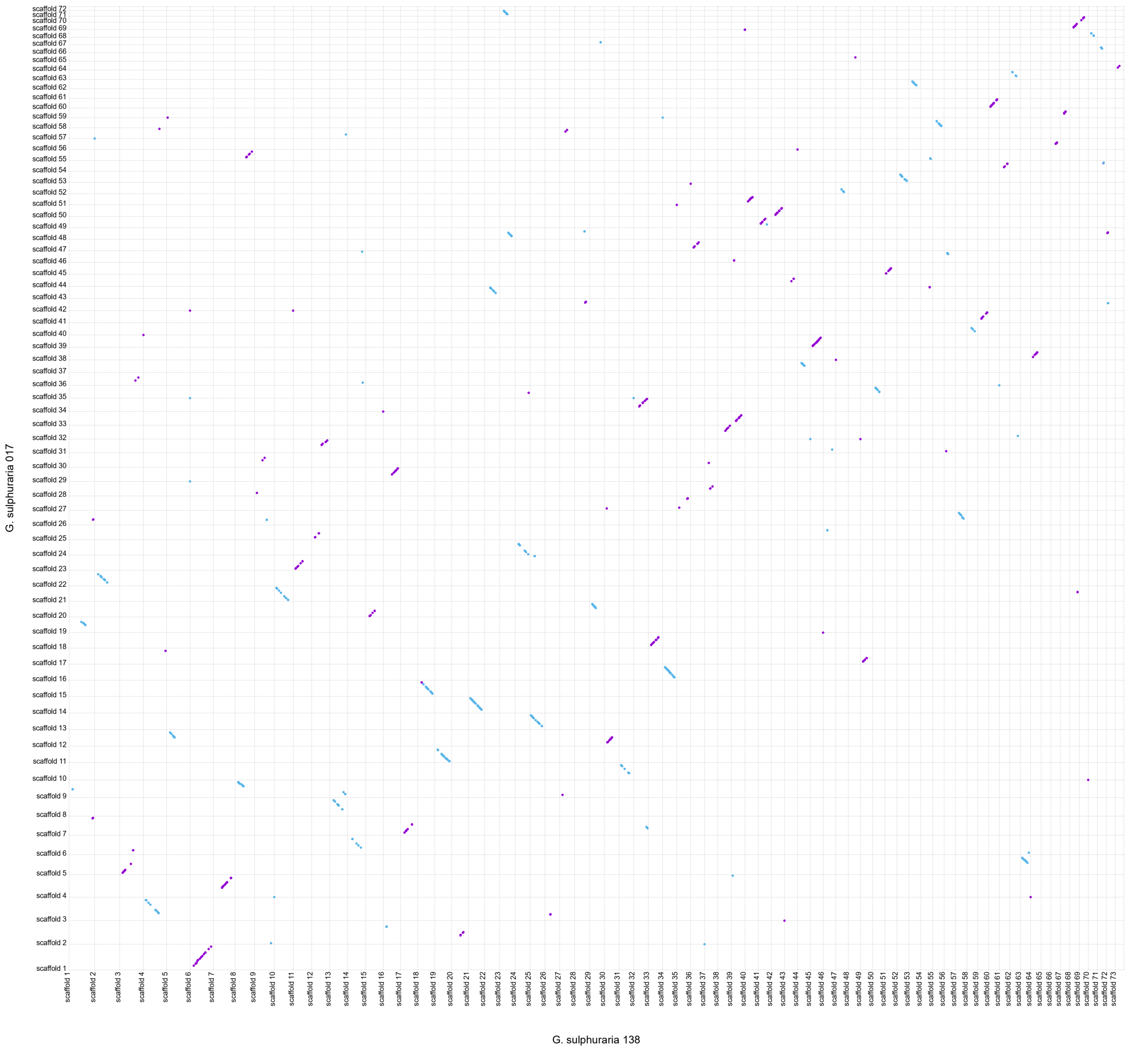

b)

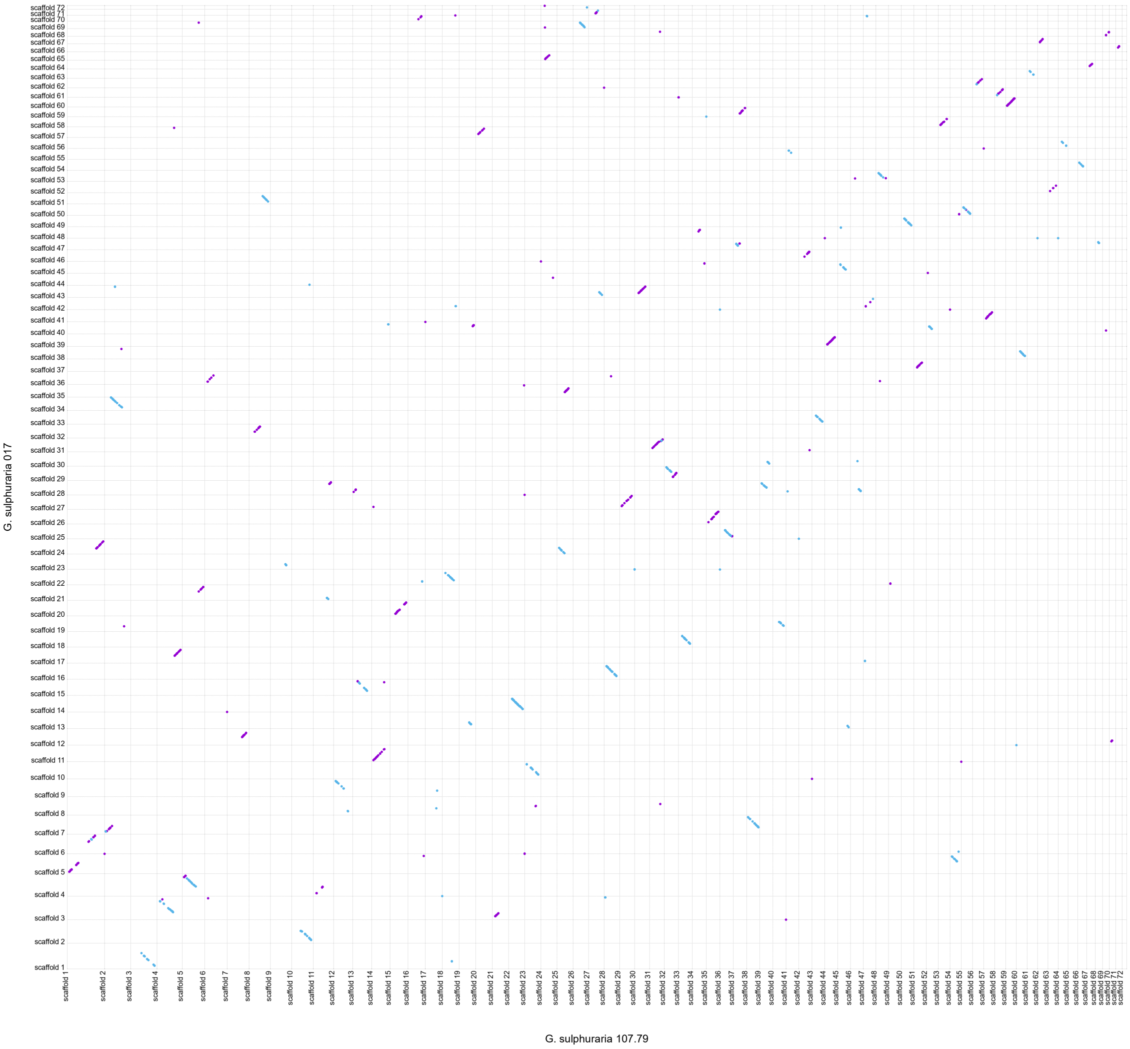

c)

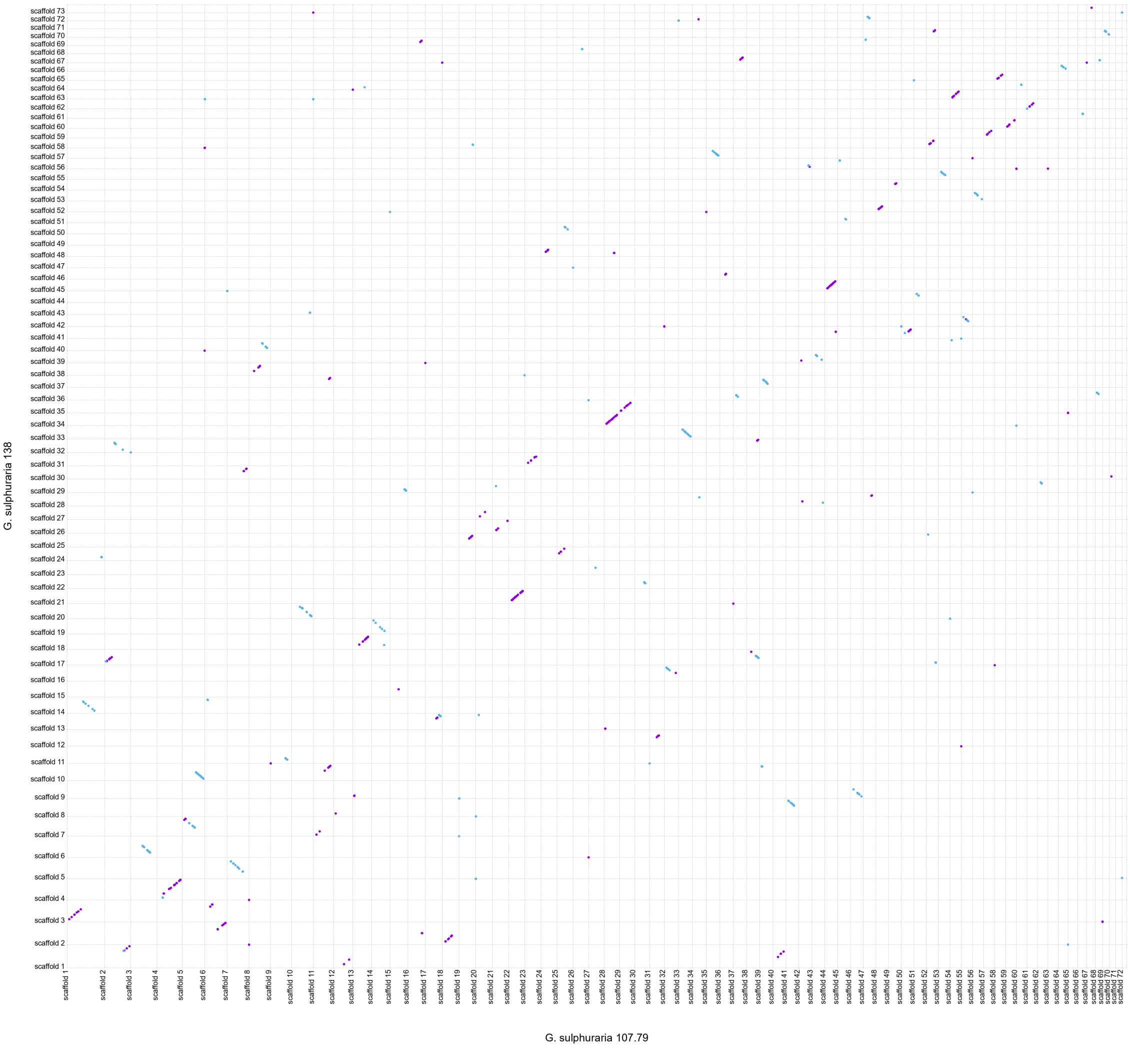

### 4. <u>All vs All Chromosome Alignment of *G. sulphuraria* SAG 107 .79</u>

All vs All *G. sulphuraria* SAG 107.79 chromosome alignment generated in nucmer, and filtered to remove alignments <100nt [1]. Forward strand alignments are shown in purple and reverse strand alignments are shown in blue.

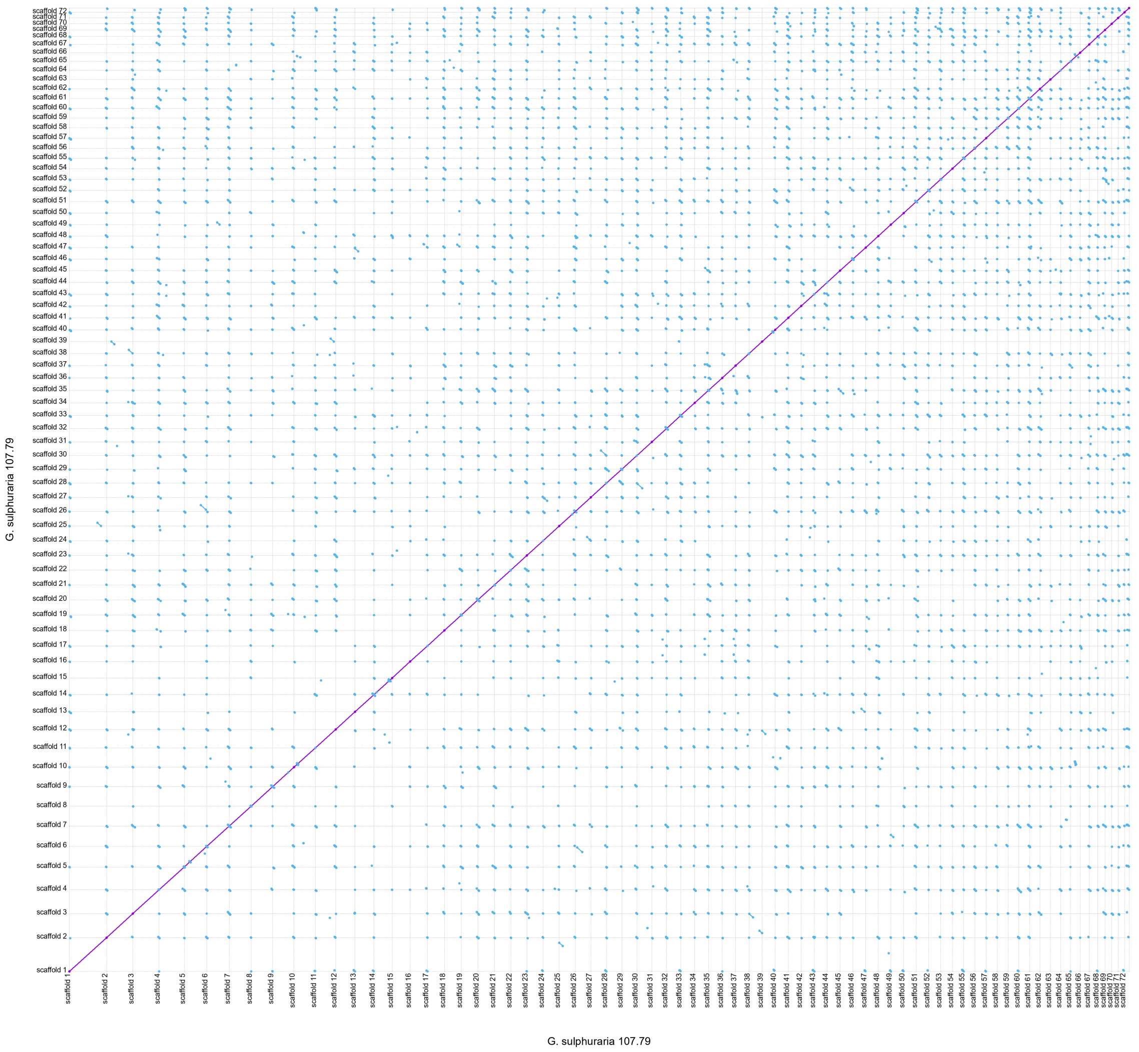

### 5. <u>Illumina Read Alignments Indicating *G. sulphuraria* is Diploid</u>

*G. sulphuraria* Illumina read alignments to the completed SAG 107.79 assembly. A) The original SAG 107.79 sequencing reads as used for polishing the assembly, b) SAG 107.79 “parent” colony taken before a 6 month single colony propagation experiment, and c) SAG 107.79 “final colony” taken at the end of the 6 month single colony propagation experiment. Reads were aligned to reference using the Burrows-Wheel Aligner [2], filtered using SAMtools [3] for properly paired reads with MQ>40, and visualised in the Integrative Genomics Viewer [4].

The section of the genome visualised, scaffold_23:47,392-48,070, shows heterozygous sites indicated by coloured bands. The first heterozygous site remains heterozygous in the final colony but the other sites are all homozygous, indicating that a recombination event in this region, followed by loss of the opposite haplotype, has occurred.

a)
b)
c)

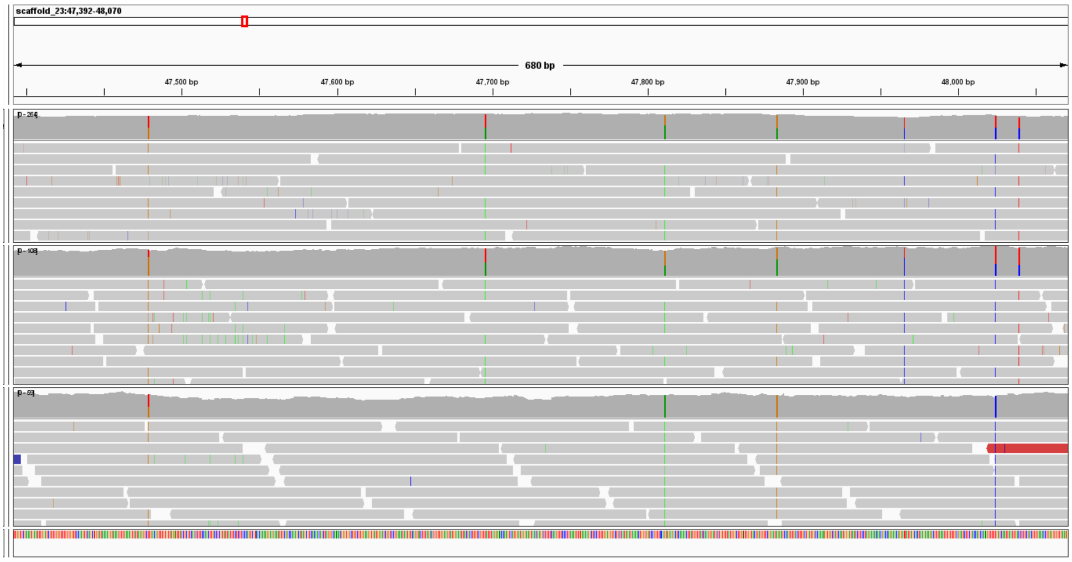
